# 22kHz and 55kHz ultrasonic vocalizations differentially influence neural and behavioral outcomes: Implications for modeling anxiety via auditory stimuli in the rat

**DOI:** 10.1101/395772

**Authors:** Camila Demaestri, Heather C. Brenhouse, Jennifer A. Honeycutt

**Affiliations:** Developmental Neuropsychobiology Lab, Northeastern University, Boston, MA 02115

**Keywords:** ultrasonic vocalizations, basolateral amygdala, anxiety, BNST, c-Fos, behavior

## Abstract

The communicative role of ultrasonic vocalizations (USVs) in rodents is well established, with distinct USVs indicative of different affective states. USVs in the 22kHz range are typically emitted by adult rats when in anxiety-or fear-provoking situations (e.g. predator odor, social defeat), while 55kHz range USVs are emitted in appetitive situations (e.g., play, anticipation of reward). Previous work indicates that USVs (real-time and playback) can effectively communicate these affective states and influence changes in behavior and neural activity of the receiver. Changes in cFos activation following 22kHz USVs have been seen in cortical and limbic regions involved in anxiety, including the basolateral amygdala (BLA). However, it is unknown how USV playback influences cFos activity within the basal nucleus of the stria terminalis (BNST), a region also thought to be critical in processing anxiety-related information. The present work sought to characterize distinct behavioral, physiological, and neural responses in rats presented with aversive (22kHz) compared to appetitive (55kHz) USVs or silence. Our findings show that rats exposed to 22kHz USVs: 1) engage in anxiety-like behaviors in the open field and elevated zero maze, and 2) show distinct patterns of cFos activation within the BLA and BNST that contrast those seen in 55kHz playback. Specifically, 22kHz USVs increased cFos density in the anterodorsal nuclei, while 55kHz playback increased cFos in the oval nucleus of the BNST. These results provide important groundwork for leveraging ethologically-relevant stimuli in the rat to improve our understanding of anxiety-related responses in both typical and pathological populations.

## Introduction

The ability to communicate cues of threat and safety is essential to the survival and proliferation of virtually all species. In humans, both verbal and non-verbal (i.e. facial expressions) cues can convey a wide range of nuanced emotional states, and it has been shown that viewing emotionally-charged facial expressions induce similar states in observers (e.g., Gump and Kulik, 1997; Wild et al., 2001). While many species do not have the capability to transfer information via intricate facial expression like humans, many produce vocalizations at distinct frequency ranges that can signal information to conspecifics. Indeed, such vocalizations can convey meaning ranging from appetitive to aversive and have been particularly well-characterized in rodents (Knutson et al., 2002; Brudzynski, 2005; Burgdorf et al., 2008; Brudzynski, 2009; Wöhr and Schwarting, 2013; Brudzynski, 2013). Rat vocalizations span ultrasonic frequencies (ultrasonic vocalizations; USVs) and indicate affective states such as rewarding pro-social states via emission of USVs in the 55kHz range (Seffer et al., 2014; Kisko et al., 2017), and fear or anxiety via emission of USVs in the 22kHz range (Brudzynski and Chiu, 1995; Kim et al., 2010). Specifically, rats emit 22kHz USVs across a variety of fearful and anxiety-provoking situations, some of which include: restraint stress (Reed et al., 2013), swim stress (Drugan et al., 2013), chronic variable stress (Mällo et al., 2009), predator odor/exposure (Blanchard et al., 1991; Fendt et al., 2018), social defeat (Kaltwasser, 1990; Tornatzky and Miczek, 1994; Kroes et al., 2007), drug withdrawal (Vivian et al., 1994; Covington and Miczek, 2003; Williams et al., 2012; Berger et al., 2013), and footshock (De Vry et al., 1993; Wöhr et al., 2005). Indeed, 22kHz USV emissions are thought to serve as alarm calls capable of warning conspecifics of possible danger and/or aversive situations (Blanchard et al., 1991; Litvin et al., 2007).

Previous work elucidating the role of natural USVs – as well as recorded USV playback – on neural and behavioral outcomes in exposed rats indicate a role for USVs in inducing affective states of fear and/or anxiety (Kim et al., 2010; Schwarting and Wöhr, 2012; Wöhr and Schwarting, 2013; Brudzynski, 2013; Briefer et al., 2018). Interestingly, rats that display high levels of anxiety-like behavior in an elevated plus maze (EPM) elicit more 22kHz USVs than those exhibiting less anxiety-like behavior (Borta et al., 2006). This affective state can be transferred via USVs to listening rat, as observed by Saito and colleagues (2016) where positive or negative cognitive bias was induced via presentation of 50kHz or 22kHz USVs (respectively) in rats in an ambiguous stimulus discrimination task. Additionally, rats exposed to 22kHz USVs also show a subsequently enhanced acoustic startle reflex (Inagaki and Ushida, 2017), further suggesting transmission of an anxiety-like behavioral state via auditory playback. While much research has been conducted examining USV generation in behavioral paradigms designed to assess anxiety (e.g., Sales, 1991; Borta et al., 2006), little attention has been explicitly paid to the impact of USV playback on subsequent behaviors in these and similar tasks. Notably, to the best of our knowledge there have been no experimental reports directly assessing the behavioral impact of USV playback in the elevated zero maze (EZM), and limited reports with regard to the open field task/arena (OFT; Endres et al., 2007; Sadananda et al., 2008). Therefore, additional playback studies are warranted to determine whether 22kHz USV playback is capable of inducing acute anxiety-like responses in rats – and to directly compare outcomes to 55kHz playback – across multiple behavioral assays.

Studies employing playback of USVs have revealed changes in brain activity in key brain regions associated with anxiety. These studies have reported significant increases in neural activation (assessed via cFos) in brain regions important for emotional valence, including the basolateral amygdala (BLA) periaqueductal grey, and the hippocampus (Sadananda et al., 2008; Ouda et al., 2016). Furthermore, research by Parsana and colleagues (2012) show that amygdala firing patterns differ depending on the type of USV playback, with an increase in firing associated with 22kHz USVs, and a tonic decrease in firing associated with 50-55kHz USVs. These findings are in line with those detailing cFos expression (Sadananda et al., 2008) and increased dopamine release (Willuhn et al., 2014) in the nucleus accumbens following 50-55kHz playback, further supporting the role of 50kHz range USVs as appetitive and socially rewarding stimuli. Interestingly, while there have been exhaustive studies looking at cFos activity following USV playback across multiple brain regions (i.e. Sadananda et al., 2008), there have been no reports on the effect of USVs on neural activation within the basal nucleus of the stria terminalis (BNST), a region that has been implicated in anxiety generation and suppression (Kim et al., 2013). Here, we expand upon the literature to determine the neural effects of both 22kHz and 55kHz USV playback on BNST cFos activity to more fully characterize their ability to induce affective states. In particular, differences in this region may serve an anxiety-specific role – and different stimuli may be dissociable based on their activation of specific BNST nuclei.

The ability for USVs to induce both behavioral and neural changes that parallel their social meaning suggest that playback may be a highly useful tool in rodent research. Indeed, it is likely that experimental presentation of a rat-generated 22kHz USV can produce a state of anxiety in a listening animal, thereby providing a model of acute anxiety-like neural and behavioral responses which can be leveraged as a translational rodent model (Jelen et al., 2003; Wöhr and Schwarting, 2013). Specifically, the playback of 22kHz USVs could be leveraged to study anxiety responsivity/salience to aversive stimuli in pathological models via acute induction of an anxiety-like state. Therefore, the present set of experiments sought to further characterize the impact of recorded USV playback in control adults on: 1) anxiety-like behavior in common behavioral assays (EZM and OFT); 2) physiological changes in heart rate during playback; and 3) changes in neural activation in brain regions associated with anxiety. Here, we provide evidence suggesting that playback of 22kHz USVs can induce increased anxiety-like behavior, changes in neural activation, and subtle alterations in physiological responses. These data support the role of 22kHz USVs as a mechanism of inducing a transient state of anxiety in rats.

## Materials & Methods

### Animals

Throughout all procedures animals were pair-housed with a same sex littermate in a temperature-and humidity-controlled vivarium (22 ± 2°C), with a 12-hour light/dark cycle (lights on at 0700h) with *ad libitum* access to food and water. Unless otherwise specified within the sections for each separate experiment below, all animals were left undisturbed in their cages except for normal husbandry procedures when not actively being run in an experiment. All experiments were performed in accordance with the 1996 Guide for the Care and Use of Laboratory Animals (NIH) with approval from Northeastern University’s Institutional Animal Care and Use Committee.

### Ultrasonic Vocalization Acquisition & Playback

The 22kHz ultrasonic vocalizations (USVs) were recorded from an adult male rat using a condenser ultrasonic microphone (Avisoft-Bioacoustics CM16/CMPA; frequency range 2kHz-200kHz) and analyzed using Avisoft-RECORDER USGH Bioacoustics Recording software (Glienick, Germany). To induce 22kHz vocalizations, the emitting rat was restrained in a DecapiCone (Braintree Scientific) and placed in a cage infused with cat urine odor inside a sound-attenuated box with a conspecific rat anesthetized with 0.3 mL/kg ketamine/xylazine to provide an audience effect (Seagraves et al., 2016). The 22kHz USV recording was high-pass filtered with a cut-off frequency of 15kHz and low-pass filtered with a cut-off frequency of 40kHz to reduce the presence of low and high frequency background noise for experimental playback. The 55kHz USV file used for playback was generously provided to us from Dr. Markus Wöhr from Philipps University (Marburg, Germany). The ultrasonic vocalizations were played back using Avisoft-Bioacoustics portable ultrasonic power amplifier (#70101) and Avisoft-Bioacoustics ultrasonic dynamic speaker (#60108) with a frequency range of ±12dB: 1-120kHz on loop for the duration of the tests described below. Representative spectrograms of the two stimuli can be seen in Figure 1.

**Fig. 1.**
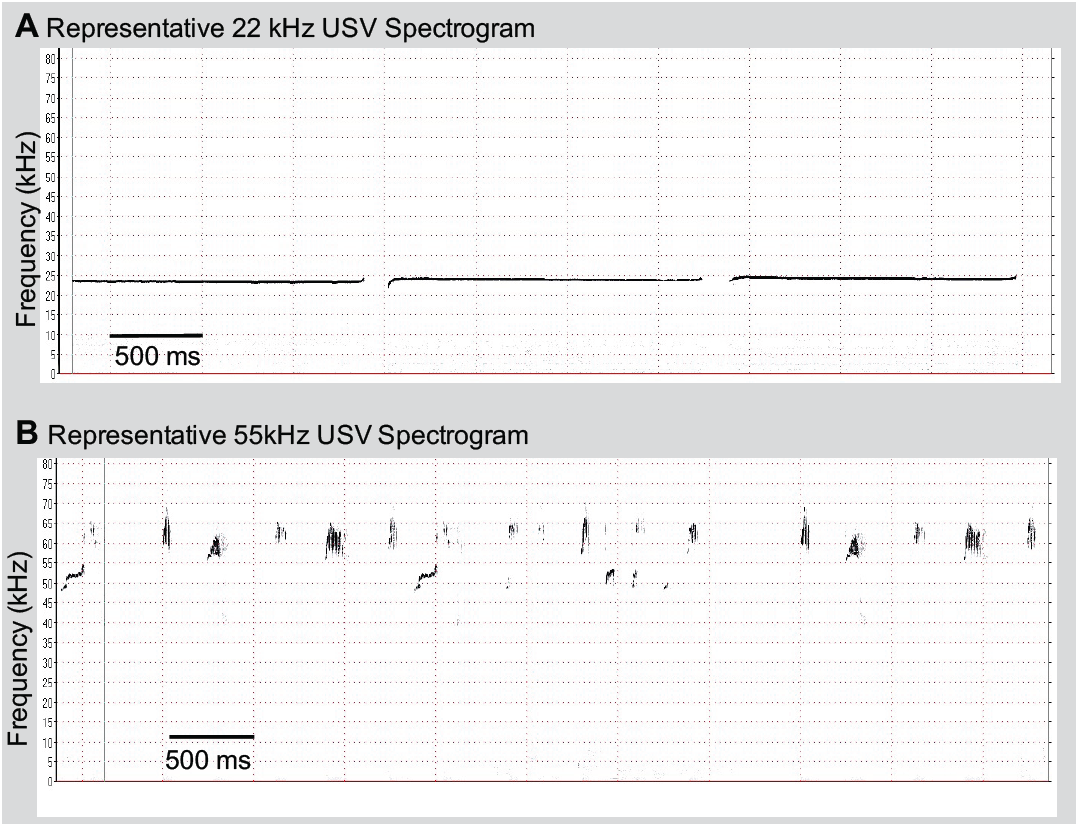
Representative spectrograms of **A**) 22kHz USV playback, and **B**) 55kHz USV playback.

### Experiment 1: USV-Evoked Changes in Open Field and cFos Expression in cFos-eGFP Rats

#### Animals & Breeding

Transgenic cFos GFP Long Evans (LE) rats, with the same transgene as the cFos-GFP mice previously described in Barth et al. (2004), were used in Experiment 1. The cFos promotor and the first 4 exons of the cFos transcript were translationally-fused to enhanced green fluorescent protein (eGFP) followed by bovine growth hormone polyadenylation signal for sequence termination. A unique advantage of this transgenic rat model is that the GFP signal can be successfully used to identify which neurons were activated during a stimulus presentation. Two hemizygotic LE-Th(cFos-eGFP)2Ottc breeding pairs were purchased from the Rat Resource and Research Center (RRRC; strain #00766; NIDA breeding program) and bred in-house via opposite-sex pair housing for one week to guarantee successful mating. Approximately 3 weeks later, both dams gave birth to litters, with date of birth marked Postnatal Day (PD) 0, and pups and dams were left undisturbed in their cages until offspring genotyping. At PD15, one ∼2mm ear punch per pup was collected, with the ear puncher sterilized with 100% EtOH between samples to avoid cross-contamination. Samples were then sent for genotyping (Transnetyx, Inc.) to confirm the presence of the transgene.

Rats positive for cFos-eGFP (*n*=14) were weaned at PD21, housed in same-sex pairs, and left undisturbed until PD60-70, when they were paired with mates from a different litter to make seven new breeding pairs. Rats were bred as described above, with ear punches from subsequent offspring also sent out for genotyping. Offspring positive for cFos-eGFP from these litters were weaned at PD21 and housed in same-sex mixed-litter pairs. For the present study, only male offspring were utilized, with female offspring dedicated to other ongoing studies. Except for normal husbandry, rats were left undisturbed until testing in adulthood (approx. PD100). A total of eighteen male LE transgenic rats were used in Experiment 1 for the Open Field Test (OFT) and subsequent immunohistochemistry analysis, as described below.

#### Open Field Test (OFT)

For Experiment 1, 18 adult male cFos-eGFP transgenic rats were randomly assigned to one of three stimulus groups: silence, 55kHz USV, or 22kHz USV (*n*=6/group). On testing day, rats were individually transported into a dimly lit testing room and left undisturbed for 5 minutes to acclimate to the testing environment. For each test, the rat was placed in the same corner of a 100cm x 100cm open field arena facing the wall. Behavior was recorded for 20 minutes via CCTV while the rats were presented with either silence, 55kHz, or 22kHz USV playback for the duration of testing via an elevated ultrasonic speaker (Avisoft Bioacoustics) approximately 30cm away from the arena. Videos were analyzed using EthoVision 9.0 software (Noldus) to determine overall distance traveled (cm), velocity (cm/s), time spent in center (s), and frequency of visits to the 40cm x 40cm center zone. Following testing, rats were returned to their home cage for one hour before euthanasia and tissue collection.

In order to account for the novelty of the acoustic stimuli, the first 10 minutes of stimulus presentation in the OFT was used to assess the initial reaction to the USV presentation (or silence), and the last 10 minutes was used to assess the final reaction to the no longer novel USV presentation (or silence). All dependent measures were subjected to Grubbs’ outlier test, which revealed no significant outliers and thus the findings for all animals are reported.

#### Tissue Collection and Immunohistochemistry

##### Perfusion & Tissue Collection

One hour following the end of USV presentation in the OFT, rats were euthanized with CO_2_ and transcardially perfused with ice-cold physiological (0.9%) saline followed by ice-cold 4% paraformaldehyde fixative solution in 0.1M phosphate buffer with a pH of 7.4. Once fixed, rats were decapitated, and brains were collected, stored in 4% paraformaldehyde solution, and allowed to post-fix for 4 days at 4°C. Brains were then switched to 30% sucrose solution in 0.1M phosphate buffered saline (PBS; pH 7.4) for cryoprotection at 4°C until ready to section. All brains were sectioned on a freezing microtome (Leica) into 40μm serial coronal sections through the entirety of the brain (excluding olfactory bulbs and cerebellum) and stored in freezing solution at -20°C until immunohistochemical processing.

##### Immunohistochemistry

To amplify and increase longevity of the GFP signal in cFos tagged cells of the cFos-eGFP brains, one series of sections from each region of interest (BLA, AU, BNST; each section spaced ∼240μm apart) were subjected to GFP immunohistochemistry. Free-floating sections were washed 3×15 min in 1× PBS and incubated with blocking buffer (0.3% TritonX-100 PBS with 3% normal donkey serum) for 1.5 hours. They were then incubated in chicken-anti-GFP igY (GFP-1020, Aveslabs) diluted 1:1000 in blocking buffer for ∼12 hours. The sections were washed 3×15 min in 1× PBS and incubated in donkey anti-chicken Alexa Fluor488 conjugate (703-545-155, Jackson ImmunoResearch) for 8 hours, followed by a 3×15 min wash in 1× PBS.

For better visualization of the regions of interest, a fluorescent NISSL stain step was added following cFos staining. Sections were incubated in NeuroTrace 530/615 red fluorescent NISSL stain (Invitrogen, N21482) diluted 1:200 in 1× PBS for one hour and then washed 3×15 min in 1× PBS. All aforementioned procedures were done at room temperature. Six slices per region of interest were mounted and coverslipped with ProLong Gold antifade reagent (Invitrogen, P36930) for later microscopy.

#### cFos Imaging and Quantification

Stained sections containing BLA (Bregma -1.72mm to -3.36mm), AU (Bregma -3.48mm to -6.00mm), and BNST (Bregma +0.48mm to - 1.32mm) were imaged at 10x (BLA, AU) or 4x (BNST) magnification using Keyence All-in-One Fluorescence Microscope BZ-X710 (Keyence Corporation of America, USA). Three sections from each region of interest where imaged bilaterally (i.e. 6 images per region; 18 images per animal). Each photomicrograph was analyzed in ImageJ (NIH) for cFos quantification. All images were subjected to the same thresholding and particle analysis parameters (i.e. size and circularity) to ensure consistent quantification across subjects/regions. The number of cFos positive cells in each image was divided by the area of the counted region to determine cFos density. All results were subjected to Grubbs’ outlier test, which revealed one significant outlier in BNST anterodorsal density in the 55kHz stimulus group. Nissl-guided delineation of regions of interest were done (with reference to Paxinos and Watson, 1997) to trace each region of interest including the BLA and the BNST, with specific BNST nuclei quantified including the adjacent anterodorsal (anterolateral and anteromedial regions), oval, and fusiform nuclei.

In addition to measuring cFos density in the BLA and BNST, we also examined the average quantification area for each region within each group to control for any discrepancies when outlining these regions for analysis (which would result in inaccurate density measures). This was not necessary for the AU because the entire (i.e. all layers) primary AU was visible in each 10x magnification image. No significant differences were observed in area for either of these regions across groups, confirming consistency in quantification across subjects (data shown in Supplementary Figure 1).

#### Statistical Analyses

All statistical analyses were conducted using Prism 7 (Graphpad Software) statistical software.

##### OFT

A two-way matching ANOVA (stimulus x time) was performed to test the main effect of stimulus (silence, 55kHz, 22kHz) on each behavior as well as any change in behavior between the first half and the second half of the OFT. Significant effects were followed up where appropriate with post-hoc Bonferroni’s multiple comparisons tests to determine group differences.

##### cFos Expression

A one-way ANOVA was performed to test main effect of stimulus on cFos density for each region of interest (BLA, AU, and BNST). A two-way ANOVA (hemisphere x stimulus) was performed to test effect of stimulus on left AU compared to right AU. Significant effects were followed up where appropriate with post-hoc Bonferroni’s multiple comparisons to determine group differences. Additional one-way ANOVAs were performed to rule out differences in BLA and BNST traced region area between groups.

### Experiment 2: USV-Evoked Changes in Elevated Zero Maze

#### Animals

Twenty-four adult male LE rats (approx. 350-450g) were purchased from Charles River Laboratories (Wilmington, MA). Upon arrival, rats were pair-housed and left undisturbed for one week to acclimate to the animal colony before the experiment began.

#### Elevated Zero Maze (EZM)

Adult male LE rats were randomly assigned to one of three groups: silence, 55kHz or 22kHz USV (n=8/group). EZM was performed on two consecutive days with four rats from each group run on each day with all groups counterbalanced for time of day (with all testing occurring between 0800h and 1300h). Rats were individually transported into a dimly lit testing room in a clean transport cage and left undisturbed for 5 minutes to acclimate to the testing environment. For each test, rats were placed in the same open arm region facing the closed arm region. Behavior was recorded for 10 minutes with a CCTV camera placed directly above the EZM, with the assigned stimulus (or silence) playing for the duration of testing. Time spent (s) in the open arm, and frequency to the open arm were calculated in 5 min bins to compare the initial reaction to the stimulus with the final reaction to the stimulus. The ultrasonic speaker (Avisoft Bioacoustics) was elevated and approx. 30 cm away from the arena. Videos were analyzed by an experimenter blind to group, with entry into an open arm defined as half or more of the rat’s body is in the open arena. Following testing, rats were returned to their home cage.

#### Statistical Analyses

Statistical analyses were conducted using Prism 7 (Graphpad Software). A two-way matching ANOVA (stimulus × time) was performed to test the main effect of stimulus on each behavior as well as any change in behavior between the first 5min and the second 5min of the EZM. Significant effects were followed up where appropriate with post-hoc Bonferroni’s multiple comparisons to determine group differences.

### Experiment 3: USV-Evoked Changes in Heart-Rate

#### Animals & Habituation

Twenty-four adult male LE rats (approx. 350-450g) were purchased from Charles River Laboratories (Wilmington, MA). Upon arrival, rats were pair-housed and left undisturbed for one week to acclimate to the animal colony. In order to limit possible artifacts on heart rate variability data based on experimenter handling, and to ensure that the rats were habituated to being in a smaller holding cage for physiological recording, a handling and habituation protocol was implemented as follows: rats were handled for five minutes/day for 7 days to habituate to experimenter contact, which was followed by two days of 5min habituation in the clear plastic recording chamber, and one day of 10min of habituation to the recording chamber, prior to testing day.

#### Electrocardiogram (ECG) Recording and Analysis

Electrocardiograms (ECGs) were recorded non-invasively using a customized ECGenie apparatus (Mouse Specifics, Inc). To eliminate circadian influences, all ECGs were recorded between 0900h and 1200h over two consecutive days. The ECGenie apparatus was used to record cardiac electrical signals at 2kHz in awake behaving rats by placing the animal in a 20cm x 15cm x 11cm recording chamber that acquires signals through footpad electrodes located on the floor of the box and transmits them to a computer for analysis.

Rats were randomly assigned to one of three groups: silence, 55kHz, or 22kHz USV playback (*n*=8/group). On test day, each rat was individually transported to the testing room in the ECG recording box and was placed 30cm away from the playback speaker and left undisturbed for a three-minute acclimation period. Following acclimation, ECG signals were recorded during a three-minute baseline period followed by a three-minute test period where they were presented with their assigned stimulus. The recording chamber was cleaned with 40% ethanol solution between each animal. Raw ECG signals were recorded using eMOUSE software (Mouse Specifics, Inc.) and analyzed as reported previously (Chu et al., 2001). ECG signals can be summarized into heart beats, each of which can be segmented into smaller standard waves, depicted as Q, R, S, T. The signal of primary interest was heartrate (HR; BPM) during baseline and test. However, additional cardiac rhythm measures were also analyzed. A representative diagram of normal ECG signals can be seen in Supplemental Figure 2A. Additional parameters analyzed were the time (ms) between subsequent R signals (time between each ventricular blood ejection), QRS complex (ventricular depolarization), ST segment (plateau phase) and QT (total duration of ventricular depolarization and repolarization, and is inversely related to heartrate) which can also be seen in Supplemental Figure 2B-E.

**Fig. 2.**
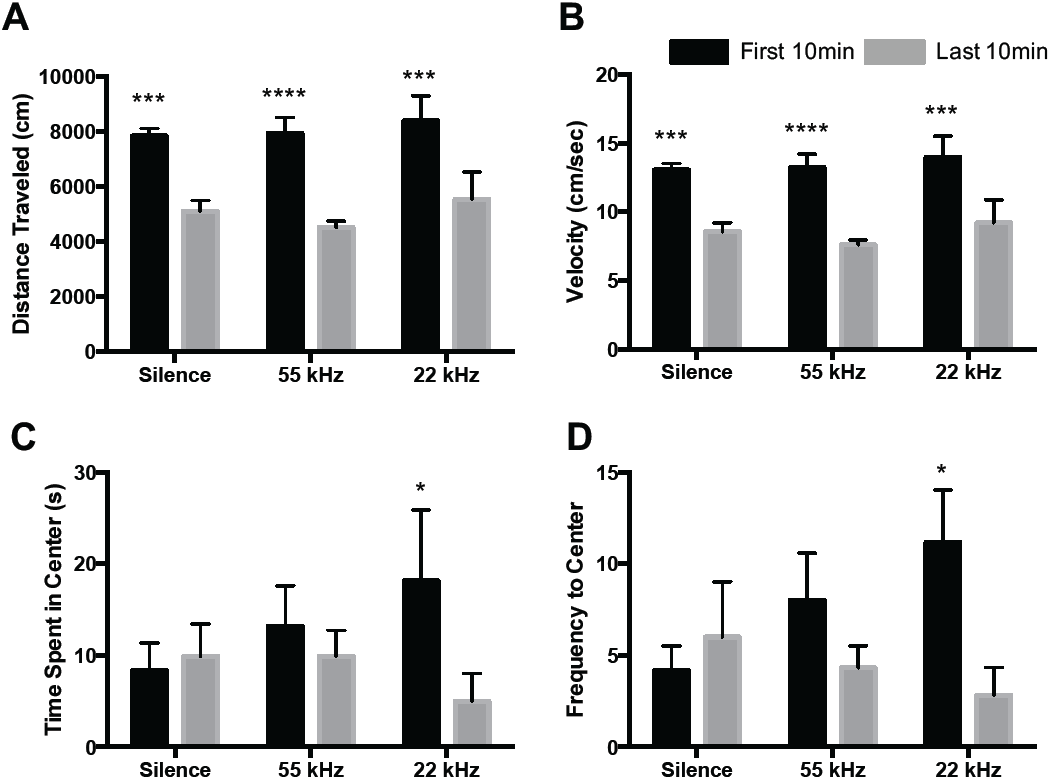
Analysis of anxiety-like behaviors in OFT in response to silence, 22kHz or 55kHz USV playback. **A**) Distance traveled (cm) decreased from the beginning of the OFT to the end of the OFT in every group. **B**) Velocity (distance traveled/1200s) decreased from the beginning of the OFT to the end of the OFT in every group. **C**) Time spent in the center zone decreased from the beginning of the OFT to the end of the OFT in response in only the 22kHz USV playback condition. **D**) Frequency to the center zone decreased from the beginning of the OFT to the end of the OFT in only the 22kHz USV condition. Data represent means ± SEM, n=6/group. \**p*<0.05, *** *p<*0.0005, **** *p*<0.0001

#### Behavioral Analysis during ECG recordings

In addition to recording ECG signals during baseline and test periods while in the ECG recording box, behavior was recorded via CCTV for later analysis by an experimenter blind to condition. Time spent mobile in the box (index of locomotion) and time spent attending to the stimulus were recorded. Time spent mobile (s) in the box was characterized by the total time spent actively investigating (i.e. general body movement, sniffing, interacting with the enclosure, etc.). Attendance to the stimulus was assessed via the difference in time spent (s) facing the direction of the USV speaker during baseline vs. test playback sessions.

#### Statistical Analyses

##### ECG Recordings and Behavior

A two-way matching ANOVA (stimulus x time) was used to test changes in HR, and mobility between baseline and test periods (3 min each). A one-way ANOVA was used to test the change in seconds spent orienting to the speaker. Additional ECG (RR, ST, QT, and QRS) signals were analyzed in the same way (Supplemental Figure 2B-E). All significant main effects were followed up with Bonferroni’s post-hoc analysis.

## Results

### Experiment 1

#### Open Field Test

Distance traveled (cm), velocity (cm/s), time spent in center (s), and number of center entries was calculated in two 10-minute bins to compare initial reaction to stimulus with final reaction to stimulus (silence, 55kHz, or 22kHz USV). A two-way ANOVA indicated a main effect of time (first 10min vs. last 10min of OFT) on both distance traveled (*F*_1, 15_ = 91.25, *p*<0.0001) and velocity (*F*_1,15_ = 91.53, *p*<0.0001). Post-hoc Bonferroni analyses showed a decrease in distance traveled from the first half of the OFT to the second half of the OFT in every group (*p<*0.001; Figure 2A) as well a decrease in velocity in every group (*p<*0.001; Figure 2B) suggesting habituation to the environment across all groups (Sestakova et al., 2013). The two-way ANOVA analyses also revealed a trend toward a main effect of time on both seconds spent in the center (*F*_1,15_ = 4.17, *p*=0.058; Figure 2C) as well as frequency of center entries (*F*_1,15_ = 4.25, *p*=0.057; Figure 2D). Follow-up post-hoc Bonferroni analyses showed a significant decrease exclusively within the 22kHz group in time spent in center and frequency of center entries from the first 10 minutes of open field when compared to the last 10 minutes of open field (*p*=0.02 and *p*=0.03, respectively) with no difference seen in silence or 55kHz USV playback groups.

#### cFos Density

cFos positive cells were counted in three regions of interest (BLA, AU, BNST) after a 20-min open field test where rats were subjected to stimulus playback of either silence, 55kHz, or 22kHz USVs. Representative Nissl images of each region of interest can be seen in Figure 3A-C, as well as representative cFos activation in each region in response to each stimulus (Figure 3D-L). One-way ANOVA analyses indicated a main effect of stimulus in cFos density in the BLA (*F*_2,15_ = 4.18, *p*=0.03; Figure 4A). BLA cFos+ cell density was significantly increased in rats exposed to 22kHz USVs compared to 55kHz USV playback (*p=*0.04), with a trend toward a significant increase compared to silence (*p=*0.058). No significant difference in BLA volume was observed between groups (*F*_2,15_ = 0.11, *p*=0.89; Supplemental Figure 1A). Within the AU, a one-way ANOVA revealed a main effect of stimulus on cFos density (*F*_2,15_ = 5.19, *p*=0.01; Figure 4B). Post-hoc Bonferroni analysis of AU indicated that cFos density was significantly increased in rats exposed to 22kHz USVs (*p=*0.03) and trending an increase in rats exposed to 55kHz USVs (*p=*0.06) when compared to silence. Additionally, only the left auditory cortex showed an effect of stimulus (*F*_2,30_ = 5.96, *p*=0.006), with cFos density increasing in response to 22kHz USVs when compared to silence (*p=*0.01 in left AU; *p=*0.09 in right AU), a finding which is in agreement with previous reports (Sadananda et al., 2008; Supplemental Figure 1B).

**Fig. 3.**
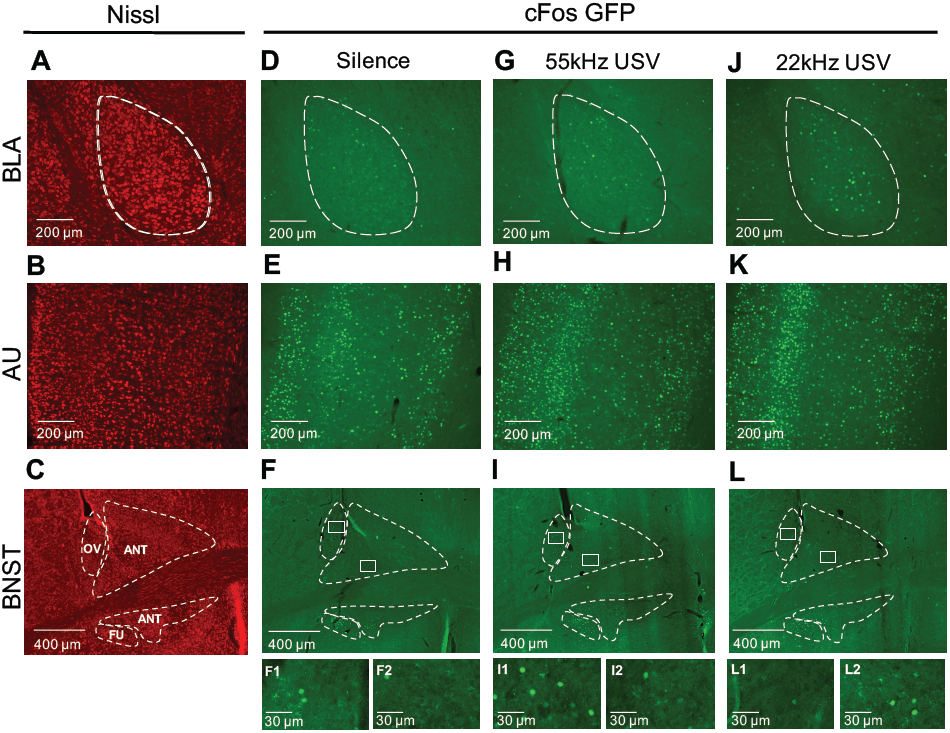
Representative images of the BLA, AU, and BNST under different auditory conditions. Nissl images and outlines of the BLA (**A**) and AU (**B**) are shown at 10x magnification, with Nissl image of BNST and nuclei (**C**) are shown at 4x magnification. To the right of these are corresponding representative images of the BLA, AU and BNST during playback of either silence (**D-F**), 55kHz USV (**G-I**), or 22kHz USV (**J-L**). Figures for the BNST include 1:1 magnified images of the oval nuclei (**F1, I1, L1**) and anterodorsal nuclei (**F2, I2, L2**) for each stimulus group. BNST Abbreviations: **OV** (oval), **ANT** (anterodorsal), **FU** (fusiform).

**Fig. 4.**
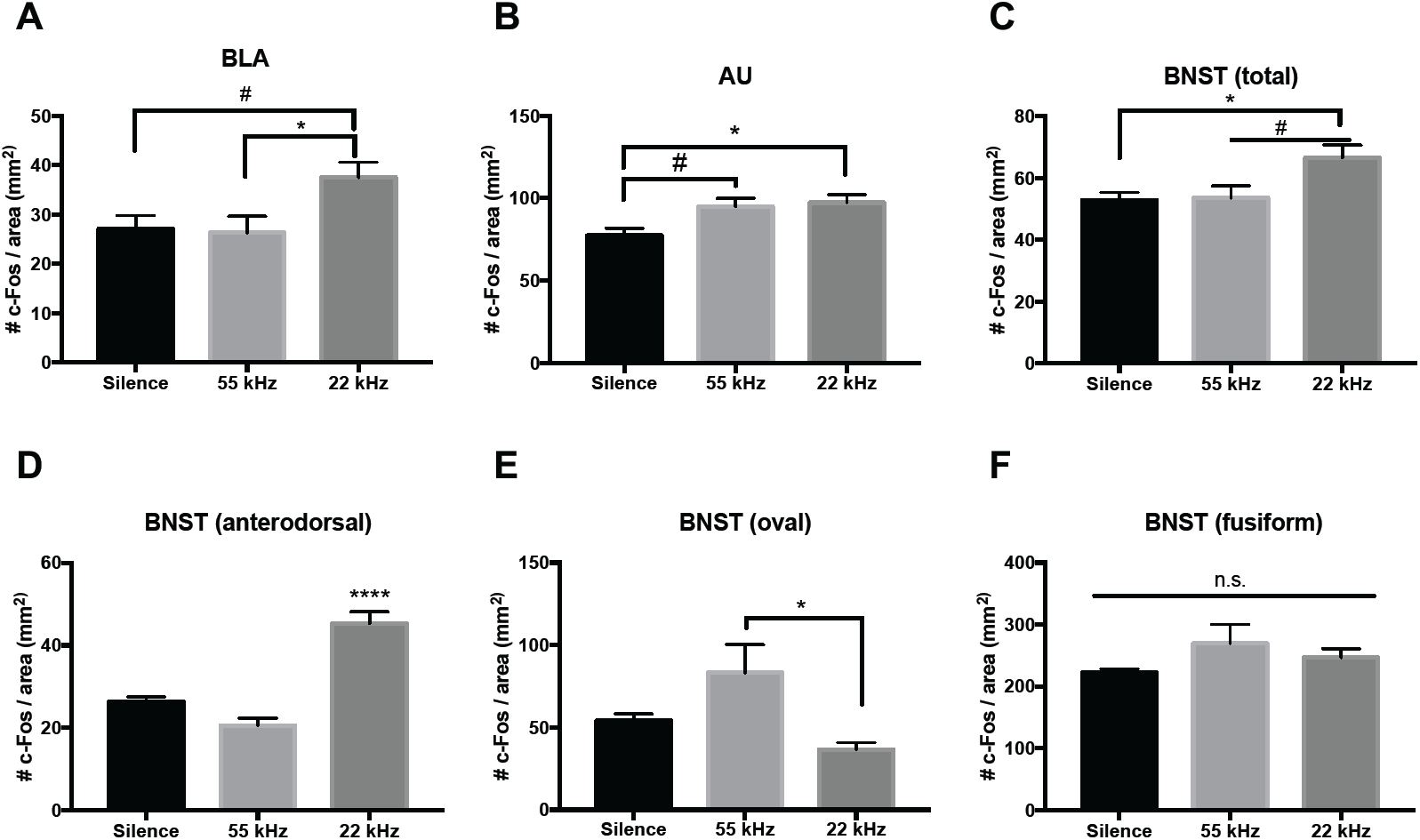
Analysis of cFos expression in the BLA, AU, and BNST in response to silence, 55kHz USV or 22kHz USV playback during a twenty-minute OFT. **A**) Density of cFos+ cells in the BLA increased when presented with 22kHz USV playback compared to both silence and 55kHZ USV playback. **B**) Density of cFos+ cells in the AU increased in response to both 22 and 55kHz USVs when compared to silence. **C**) Density of cFos+ cells in the BNST is increased in the 22kHz group. **D**) cFos+ cell density in the BNST anterodorsal nuclei is significantly increased in the 22kHz group compared to both silence and 55kHz. **E**) cFos+ cell density in the BNST oval nucleus increased in response to 55kHz compared to 22kHz USV presentation. **F**) No difference in cFos+ cell density was seen in the BNST fusiform nucleus. Data represent means ± SEM, n=6/group (n=5 in 55kHz group BNST total, and BNST anterodorsal analyses). #*p<*0.06, \**p*<0.05, \*\*\*\**p*<0.0001

One-way ANOVA analysis of cFos+ cell density in the total BNST counted revealed a significant main effect of stimulus *(F*_2,14_ = 4.837, *p*=0.025) with an increase in response to 22kHz USV playback when compared to both silence (*p=*0.035) and 55kHz USV playback (*p=*0.06; Figure 4C). Additional analysis within the BNST showed an effect of stimuli in both the anterodorsal nuclei (*F*_2,14_ = 38.33, *p*<0.0001) and oval nucleus (*F*_2,15_ = 5.143, *p*=0.019), but not the fusiform nucleus (*F*_2,15_ = 1.447, *p*=0.266; Figure 4D-F). Contrasting effects were seen in the anterodorsal nuclei and the oval nucleus. The anterodorsal nuclei showed an increase in cFos+ cell density in response to 22kHz USV playback (*p<*0.0001) when compared to both silence and 55kHz USVs, while the oval nuclei showed an increase in response to 55kHz USV playback (*p=*0.018) when compared to 22kHz USVs.

### Experiment 2

#### Elevated Zero Maze

Two-way ANOVA analyses revealed a main effect of stimulus (*F*_2,21_= 5.241, *p*=0.01) on time spent in the open area of the EZM (Figure 5A). Post-hoc analyses revealed significantly less time spent in the open arm when exposed to 22kHz USVs compared to silence (*p=*0.01). Furthermore, rats in the 22kHz USV group spent less time in open areas during the last five minutes compared to the first five minutes of the test (*p=*0.05), and significantly less time in open areas during the last five minutes when compared to the silence group (*p=*0.0005). No difference in time spent in open areas of the EZM was seen between time bins as a result of 55kHz USV playback (*p>*0.1). Additionally, no effect of stimulus (*F*_2,21_ = 0.22, *p*=0.78) or time (*F*_1,21_ = 2.58, *p*=0.12) was found in frequency to the open area of the EZM (Figure 5B).

**Figure 5:**
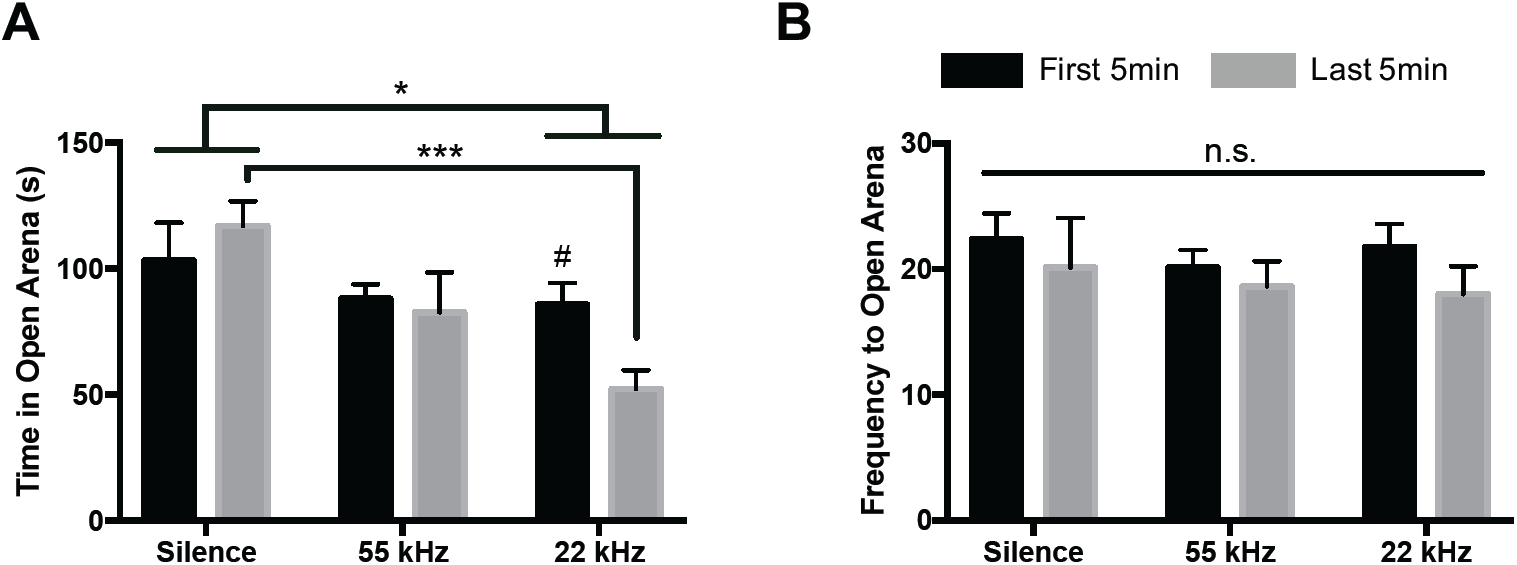
Analysis of anxiety-like behaviors during 10min EZM in response to USV playback in first vs. last 5min of testing. **A**) In general, time (s) spent in the open arena of the EZM in response to 22kHz USVs is significantly less than in response to silence. Time spent in open areas decreased from the first five minutes of the test to the last five minutes of the test in response to 22kHz USVs. **B**) Frequency to open areas of the EZM does not change as a function of time or stimulus type. Data represent means ± SEM; n=8/group. *#p*<0.06, *p<0.05, *** *p<*0.0005

### Experiment 3

#### ECG recordings & Behavior

To determine the impact of USV presentation timeline (baseline vs. stimulus presentation) and/or stimulus (silence, 55kHz, or 22kHz USV), a two-way ANOVA was conducted. On HR data, analyses revealed a main effect of presentation timeline (*F*_1,21_ = 10.20, *p*=0.004), in addition to a timeline x stimulus interaction (*F*_2,21_ = 3.82, *p*=0.03), with HR decreasing from baseline to test period in the silence group (*p=*0.001), but not in the 55kHz or 22kHz USV groups (*p>*0.99; Figure 6A). Similar patterns were detected in QT signals, with an interaction of timeline x stimulus (*F*_2,20_ = 4.68, *p*=0.02), showing an increase in time (ms) between QT signals during stimulus presentation compared to baseline in the silence group (*p=*0.01), but not in 55kHz or 22kHz USV groups (*p>*0.99; Supplemental Figure 2B). No significant effects of timeline or stimulus on ST, RR, or QRS ECG signals were identified (Supplemental Figure 2C-E).

**Figure 6:**
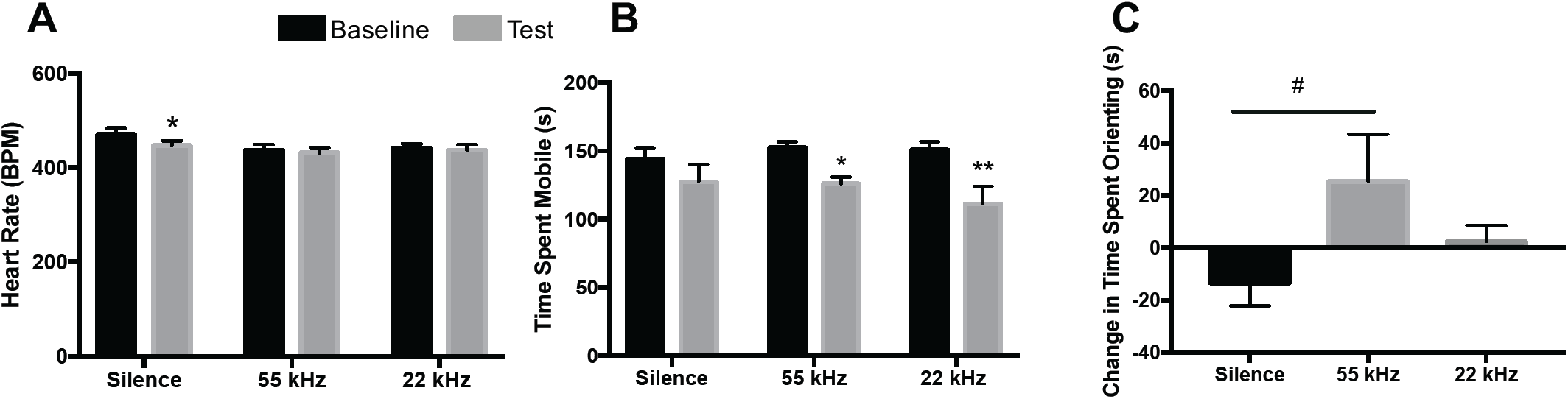
Heart rate and behavioral changes during baseline (3min) and USV playback (3min). **A**) Heart rate (BPM) decreases only in the silence group from baseline to testing, with no change in 55kHz or 22kHz groups (n=8/group). **B**) Time (s) spent actively mobile significantly decreased from baseline to test in both the 55kHz and 22kHz groups, with no change seen in the silence group (silence: n=8; 55 and 22kHz USVs: n=7/group) **C**) Difference in time (s) spent oriented toward the speaker from baseline to test/presentation (test(s)-baseline(s); n=8/group). Data represent means ± SEM. #*p*=0.06; \**p*<0.05; \*\**p<*0.01

Time spent mobile during ECG recording was monitored during both the three-minute baseline and three-minute stimulus presentation periods. Analyses show a main effect of time on mobility (*F*_1,19_ = 25.23, *p*<0.0001) within the testing apparatus (Figure 6B). Rats exposed to 55kHz USV playback and 22kHz USV playback displayed a significant decrease in mobility from baseline to test periods (*p=*0.039 and *p=*0.002, respectively). The change in time spent oriented toward the speaker during USV playback was not significantly different between stimuli, *F*_2,21_= 2.607, *p*=0.09), though multiple comparison tests revealed a modest increase in time spent oriented toward the speaker during playback of 55kHz USVs (*p=*0.06; Figure 6C).

## Discussion

### 22kHz and 55kHz USVs differentially mediate cFos in the BLA, and BNST

The present work demonstrates a role for both 22kHz and 55kHz USVs in mediating cFos density in a region-dependent manner, particularly within anxiety-related brain regions (BLA, BNST). In the BLA, a robust increase in cFos density was observed in response to 22kHz USV playback when compared to both silence and 55kHz USV playback. This finding is supported by previous work indicating that 22kHz, but not 55kHz, USV playback increases overall BLA activation compared to non-social stimuli and/or silence (Ouda et al., 2016; Sadananda et al., 2008). BLA activity has been shown to be indicative of anxiety-like states in the rat (Davis and Whalen, 2001; Vyas et al., 2004; Daskalakis et al., 2014), and therefore this change could be an index of induced anxiety in response to a socially-relevant aversive stimulus. The fact that 55kHz USVs evoked no significant changes in cFos density in comparison to rats exposed to silence further supports this interpretation. Importantly, it is unlikely that there were fundamental differences in the rats’ ability to perceive the two distinct stimuli that were driving this finding, as both 22kHz and 55kHz USVs showed comparably increased cFos density in the primary auditory cortex compared to rats exposed to no stimulus playback.

While the BLA is a well-known and highly studied locus of anxiety-related behavioral states (Tye et al., 2011; Wang et al., 2011; Janak & Tye, 2015; Perusini et al., 2015), the BNST also plays an important role in an organism’s response to anxiety-inducing stimuli (Walker et al., 2003; Kim et al., 2013; Avery et al., 2016). While it has been demonstrated that the BNST nuclei each differentially subserves its highly heterogenous anxiety-regulating functions (Dabrowska et al., 2013; Kim et al., 2013; Gungor and Paré, 2016), there are no reports detailing how the BNST responds to USV playback in the rat. Here, we provide the first report of USV-induced changes in cFos density following both 22kHz and 55kHz USV playback in adult rats. Specifically, our data shows an increase in total BNST cFos density in response to 22kHz USVs compared to both silence and 55kHz USVs; a finding that further supports the capability of 22kHz USVs to induce an anxiety-like state in a listening rat.

In order to understand how this was driven by individual nuclei within the BNST, it was subdivided into 3 regions thought to be involved in anxiety and stress, including: anterodorsal (anterolateral and anteromedial nuclei), oval nucleus, and fusiform nucleus (Daniel and Rainnie, 2016). While no significant differences were found between groups in the fusiform nucleus, the anterodorsal and oval regions showed differential activation patterns. In rats exposed to 22kHz USV playback, a significant increase in cFos density was observed within the anterodorsal region compared to both 55kHz and silent conditions. Conversely, rats exposed to 55kHz USV playback showed a significant increase within the oval nucleus when compared to 22kHz yet was no different to silence. These findings suggest that USV playback – depending on the social relevance – induce opposing patterns of neural activation within the BNST, providing insight into the different roles these nuclei play in moderating both appetitive and anxiety-evoking stimuli. Similarly, Butler and colleagues (2016) found that ferret odor induced increased cFos in the anteromedial nuclei (i.e. anterodorsal), with no changes observed in the oval (which they refer to as the lateral BNST).

Interestingly, past evidence suggests that optogenetic activation of the oval is anxiogenic, whereas activation of the anterodorsal is anxiolytic (Kim et al., 2013). However, it is important to specify that BNST subregions are largely heterogenous with regard to cell type (i.e. CRF, GABA, DA, GLU, etc), and also project to a wide range of brain regions, including the amygdala, hippocampus, hypothalamus, nucleus accumbens, VTA, etc. (Dong et al., 2001; Larriva-Sahd, 2006; Silberman and Winder, 2013; Partridge et al., 2016), thereby supporting pathways which can either mediate or exacerbate anxiety-like responses. Adding nuance to this narrative, activation of different subtypes of neurons within the BNST lead to dissociable responses, such that activation of GABAergic neurons within the BNST impart anxiolysis, whereas activation of glutamatergic projections from the BNST is anxiogenic (Jennings et al., 2013). Furthermore, within the oval, there is a high concentration of CRF neurons, which are mostly GABAergic (Dabrowska et al., 2013). Although our results do not elucidate on the cell-specific mechanisms by which USV playback is exerting its differential effect, it is possible that the differences in cFos density seen in response to 22kHz and 55kHz playback in the anterodorsal and oval nuclei may be modulated by different cell types. Due to the complexity of the BNST, the specific anxiogenic and/or anxiolytic roles of USV playback within distinct nuclei are difficult to pinpoint without additional cell-and circuit-specific analyses. However, it is clear that the BNST is distinctly sensitive to USVs in the rat, with each vocalization preferentially evoking different neural responses, which we also observed to be reflected in behavior.

### Behavioral effects of 22kHz and 55kHz USVs

The present behavioral findings provide evidence that playback of 22kHz USVs – but not 55kHz USVs – induce an acute anxiety-like state in the rat. Two behavioral paradigms (OFT and EZM) were utilized to characterize anxiety-like behaviors in rats in response to USV playback to determine the suitability of 22kHz USV stimuli to instill anxiety in control animals. Previous work suggests that 22kHz USVs decreases overall locomotion in the OFT, while 55kHz USV playback increases locomotion (Wöhr and Schwarting, 2007). However, other groups point to a lack of behavioral effects during USV presentation, with no differences observed in the OFT (Endres et al., 2007). While these prior findings stand in contrast to one another, the present OFT findings add an additional nuance, indicating that there may be a time-dependent effect of presentation. While there were no between groups differences in time spent in center or frequency to center between stimuli groups, there was a significant within-subjects decrease in both of these measures between the first and last 10 min of testing exclusively in the 22kHz group.

Though between-group changes were not seen based on stimulus, this decrease between the first and last 10 min of stimulus presentation on time in – and frequency to – center suggests that the 22kHz USVs induce a differential effect across time. While rats do not initially show an anxiogenic effect of 22kHz presentation in the OFT, it is clear that sustained presentation has the capability of inducing anxiety-like behavior. This could be attributed to a variety of factors, including initial novelty of USV playback inducing a moderate increase in time spent in center. There is also evidence suggesting that prior self-eliciting of aversive USVs may be necessary to induce anxiety-like behaviors during playback (Parsana et al., 2012), though our data and research from other groups indicates that this may not actually be needed for anxiety to be induced (Calub et al., 2018).

These data are the first to show a differential response to the rats’ initial reactions to the 22kHz stimulus compared to a later, secondary response during sustained playback. This effect was also replicated in the EZM, with rats spending less time in the open areas of the EZM during the last 5min compared to the first 5min during 22kHz playback. In contrast to the OFT, this behavioral paradigm showed an overall reduction in the open area time in response to 22kHz USVs when compared to silence, an effect which has also been documented following restraint stress or administration of an anxiogenic drug (Braun et al., 2011). The innate anxiety-provoking effect of the EZM compared to the OFT in control animals might have exacerbated the rats’ responsivity to the 22kHz USVs, making the anxiety-like responses more apparent in this assay. Importantly, this is the first report showing that 22kHz USV playback increases anxiety-like behavior in the EZM in comparison to both silence and 55kHz USVs.

### Effects of USV presentation during ECG recordings

Change in HR is modulation by the sustained regulation of the parasympathetic and sympathetic nervous systems. Under resting conditions, the vagal nerve tonically inhibits the parasympathetic nervous system, with low vagal tone associated with increased anxiety in children (El-Sheikh et al., 2001). Thus, successful modulation of the parasympathetic system via the vagal nerve significantly impacts the relationship between HR and anxiety. Here, we show that a decrease in HR from baseline to test only occurs in the group exposed to silence. This decrease in HR is likely a result of decreased physiological arousal, potentially indicating a habituation effect of being in the holding box during the experiment. HR remained unchanged in response to USVs, despite that overall mobility during playback was significantly decreased in both groups compared to baseline. The lack of change in HR in the USV groups is likely indicative of successful activation of the vagal nerve in inhibiting the sympathetic nervous system, with concurrent increased alertness to both stimuli. We hypothesize that this complex modulation would be derailed in rats that exhibit high levels of anxiety following experimental manipulations (i.e. early life adversity, chronic stress, etc.), and would therefore exhibit an increase in HR during aversive USV playback. Rats in the 55kHz condition also show a slight, but notable, increase in time spent oriented to the speaker, which is consistent with an increase in approach behavior in response to an appetitive auditory stimulus (Wöhr and Schwarting, 2007; Sadananda et al., 2008).

It must be discussed, however, that there are significant limitations that exist with HR analysis. The signals received from the ECG recorder varied extensively between animals during both baseline and test conditions. For example, the range in the number of recorded events over three minutes for each animal was 21-244. Signals are successfully recorded when the rat is on all fours, immobile for at least 3 seconds, and with nothing obstructing the grounding of the ECG electrode pad (i.e. tail movement, urine, stool). Thus, if the animal were to move extensively, groom, defecate, etc., then the signals were difficult to detect. It is likely that this variability may have influenced the ability to interpret the data, and therefore additional research should be done to determine whether USV playback is able to modulate cardiovascular output. Additionally, the decrease in HR in only the silence group from baseline to test could suggest that the initial habituation procedure was insufficient to ameliorate the change in arousal from being held in the small enclosure, which may also explain why baseline HR was ∼50BPM higher than has been previously reported for adult rats (Barnard et al., 1976), and might contribute to the lack of change in HR in the USV groups. Importantly, it is worth mentioning that these experiments were conducted in control animals, and that it is likely that rats with pathological anxiety might show differential regulation of HR.

## Conclusion

The present work provides additional evidence that aversive 22kHz USV playback is capable of inducing an acute state of anxiety in control rats, a finding which is supported by concomitant changes in the BLA and BNST. Importantly, these data are the first to describe 22kHz USV-evoked anxiety-like responses in the EZM, in addition to the first to evaluate the effect of USV playback on BNST cFos activation. Indeed, we describe a robust difference in BNST reactivity to USV playback that suggests a differential role of individual BNST nuclei in processing responses to socially-relevant aversive and appetitive auditory stimuli. The behavioral findings suggesting an increase in anxiety-like behavior in both the OFT and the EZM, paired with the changes seen in BLA and BNST activation, support the ability for 22kHz USV playback to induce an acute state of anxiety. Based on this evidence, we suggest that 22kHz USV playback is an effective and ethologically-relevant method of anxiety induction, with the potential to be leveraged to examine differential responses to anxiety in animal models of pathology.

This article has supplementary materials.

## Acknowledgments

The authors would like to thank Dr. Markus Wöhr for generously providing the 55kHz USV recordings for use in the present work, and Dr. Thomas G. Hampton of Mouse Specifics for his assistance in the modification and use of his ECGenie system for recording the ECGs for this work. Additionally, we would like to sincerely thank Shayna Peterzell, Kei Inge, Alexa Soares, and Habiba Shaheed for their assistance in collecting and coding behavioral data and Nicholas Tinsley for assistance in USV processing and filtering.

## Funding

The research described in this manuscript was supported in part by a NARSAD Young Investigator Grant from the Brain & Behavior Research Foundation (awarded to JAH), and the National Institutes of Health (5R01MH107556 awarded to HCB).

## Competing Interests

The authors have no competing interests to declare.

